# The promise of long-read RNA-seq: reducing bias in analyses of allele imbalance

**DOI:** 10.1101/2025.10.14.682301

**Authors:** Nadja Nolte, Marko Petek, Pablo Angulo Lara, Logan Mulroney, Francesco Nicassio, Fabio Marroni, Lauren McIntyre

## Abstract

Inaccurate allele and gene expression counts due to map bias and genome ambiguity lead to high false positive and false negative rates in studies of allelic imbalance. We demonstrate that long read RNA-seq and straightforward quality control measures can be used to reduce bias in allele counts in case studies from four species: *Drosophila melanogaster*, a diploid insect; *Solanum tuberosum*, an autopolyploid plant; *Pongo abelii*, a highly heterozygous diploid primate, and *Homo sapiens*. We recommend 1) mapping to a personalized genome to increase the number of allele assignments 2) tracking multimapping reads and tuning mapping parameters to ensure accurate allele and gene expression counts and 3) evaluating apparent extreme allele bias to identify errors in genome assembly and annotation. We show that these steps can be executed in a straightforward manner and recommend tools for each step.

## Introduction

Reference genomes, typically represented as a single haplotype sequence for a species (Fig. 1a), have been a transformative milestone in modern biology. The quantification of allelic expression relies on being able to separate sequencing reads from RNA-seq experiments based on genetic variation, and bias in allele counts can result in high false positive rates (Degner et al., 2009). Single Nucleotide Polymorphisms (SNPs) N-masking is a common technique to reduce bias, however incorrect assignment of reads to genes and alleles persists (Pandey et al., 2013). Alternatively, personalized genomes that incorporate allelic variants of an individual (Fig. 1b) have been shown to reduce bias in quantifying allele counts for multiple species (Adduri & Kim, 2024; Castel et al., 2015; Crowley et al., 2015; Degner et al., 2009; Demirdjian et al., 2020; Dobin et al., 2013; Glinos et al., 2022; Graze et al., 2012; Munger et al., 2014; Pandey et al., 2013; Rozowsky et al., 2011, 2023; Vaddadi et al., 2023; Wu et al., 2023).

**Figure 1:**
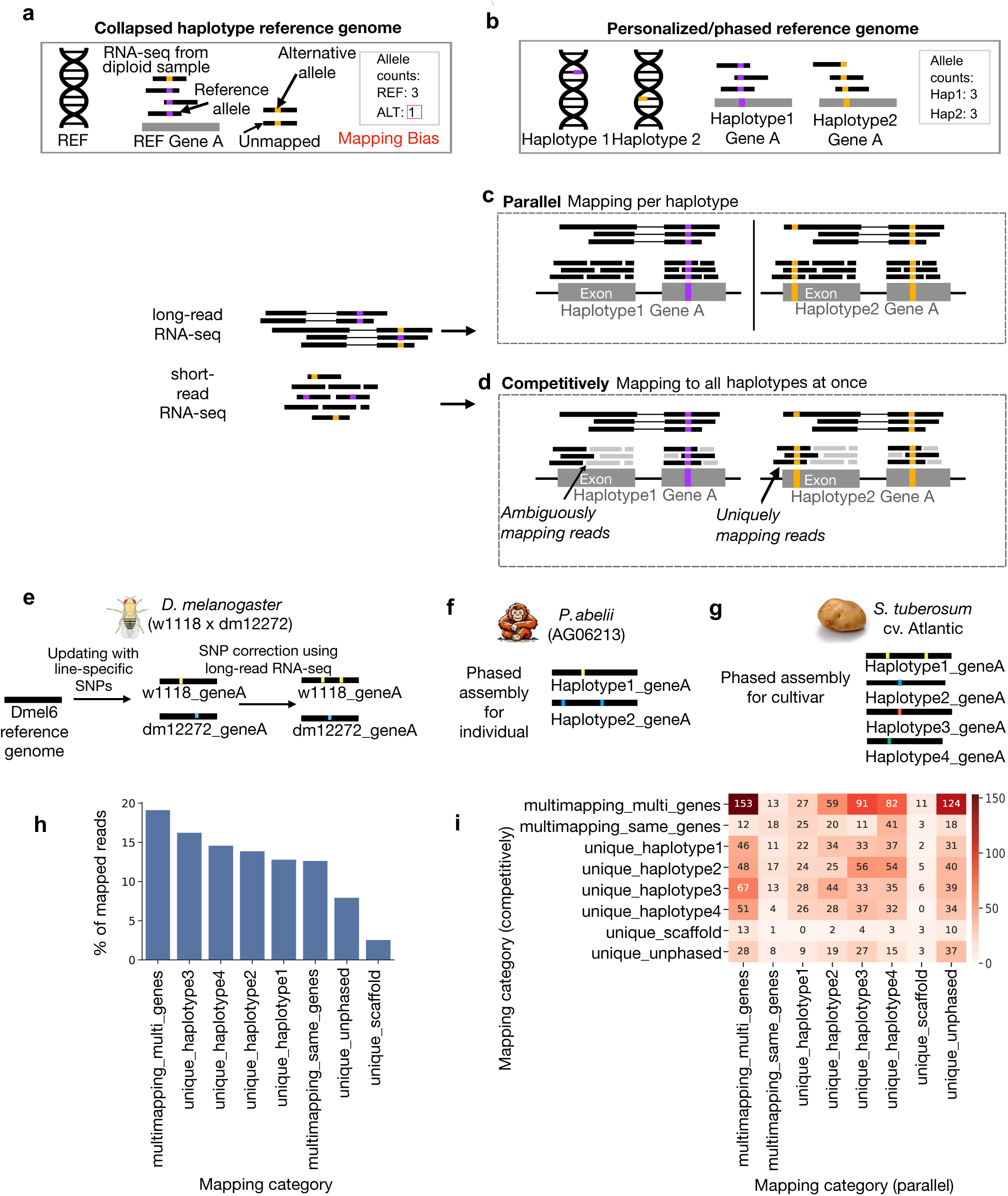
The reference used affects equivalence groups. Tracking multi-mapping results in equivalent allele and genotype counts regardless of the reference used *(a)* Haploid reference genomes and mapping bias. Reads containing the reference allele may be more likely to align than those containing alternative alleles. *(b)* Haplotype-phased personalized reference genomes mitigate mapping bias. RNA-seq data can be mapped either *(c)* in parallel to each haplotype or *(d)* to a combined genotype reference. The proportion of uniquely mapped reads (black) versus multimapping reads (grey) differs in these two strategies. Long-read RNA-seq is expected to have less ambiguity and overall fewer multi-mapping reads compared to short-read RNA-seq. *(e–g)* Case studies for the two mapping strategies applied to *(e)* diploid *D. melanogaster* using a SNP-updated personalized reference, *(f)* diploid *P. abelii* with an phased assembly for the individual assayed, and *(g)* tetraploid *S. tuberosum* with a phased assembly for the cultivar assayed. *(h)* Barplot of the number of reads in each category having the identical mapping location in separate and competitive mapping *for S. tuberosum.* Reads mapping to only one allele are assigned to mapping category “unique_haplotype[1,2,3,4]”, reads mapping equally well to multiple alleles of a gene are assigned as “multimapping_same_gene” and reads mapping equally well to multiple allele and genes are classified as “multimapping_multi_genes” *(i)* Heatmap showing number of reads with different mapping location in separate and competitive mapping *for S. tuberosum*.

The cost of personalized genomes was limiting their use. Genetic reference panels, such as the Drosophila Genetic Reference Panel (Mackay et al., 2012), the Mouse Collaborative Cross (Collaborative Cross Consortium, 2012), and The Arabidopsis Information Resource (TAIR) (Rhee, 2003), re-sequenced sets of nearly homozygous genotypes, enabling individual researchers to use stocks of sequenced genotypes. Crosses between these genotypes created heterozygotes with known allelic variation and enabled personalized genomes in studies of allelic imbalance (e.g. Churchill et al., 2012; Crowley et al., 2015; Fear et al., 2016; Graze et al., 2012; L. G. León-Novelo et al., 2014). Recent advancements in sequencing technology and genome assembly algorithms have made phased *de novo* reference genomes more accessible and affordable (Miga & Eichler, 2023; Sarashetti et al., 2024). Haplotype phased cultivar-specific genomes have been reported for agriculturally important plants, such as apple (Su et al., 2024), potato (Godec et al., 2025; Hoopes et al., 2022; Sun et al., 2022), and rice (Feng et al., 2021). Other species, such as honeybee (Bresnahan et al., 2023) and orangutan (Mao et al., 2024), have also released phased haplotype references. In humans, major efforts are underway to capture individual haplotype variation in pangenomes (Liao et al., 2023). Haplotype-resolved pangenomes reduce bias in read mapping (Sibbesen et al., 2023) and can be used to create personalized references (Vaddadi et al., 2023).

In studies of allele imbalance, unbiased estimates of alleles and gene expression are crucial to avoiding large type I error rates (Akama et al., 2014; Glinos et al., 2022; Graze et al., 2012; T. Kuo et al., 2018; L. G. León-Novelo et al., 2014; Munger et al., 2014; Page et al., 2013; Rozowsky et al., 2011). Accurate estimates of gene expression directly impact allelic imbalance, and there is a relationship between gene expression and the power to detect allelic imbalance (L. León-Novelo et al., 2018; Raghupathy et al., 2018; Sherbina et al., 2021). Discarding the multi-mapping reads can lead to biased estimates of gene expression (Deschamps-Francoeur et al., 2020; L. León-Novelo et al., 2018; Raghupathy et al., 2018; Robert & Watson, 2015; Szabelska-Beresewicz et al., 2023; Van De Geijn et al., 2015; Zytnicki, 2017) by underestimating gene expression in gene families (Robert & Watson, 2015; Szabelska-Beresewicz et al., 2023, Choi 2019), biasing functional enrichment analysis (Almeida Da Paz et al., 2024), resulting in higher levels of variability between replicates and increased error rates (Choi et al., 2019) (Fig. 2g).

**Figure 2:**
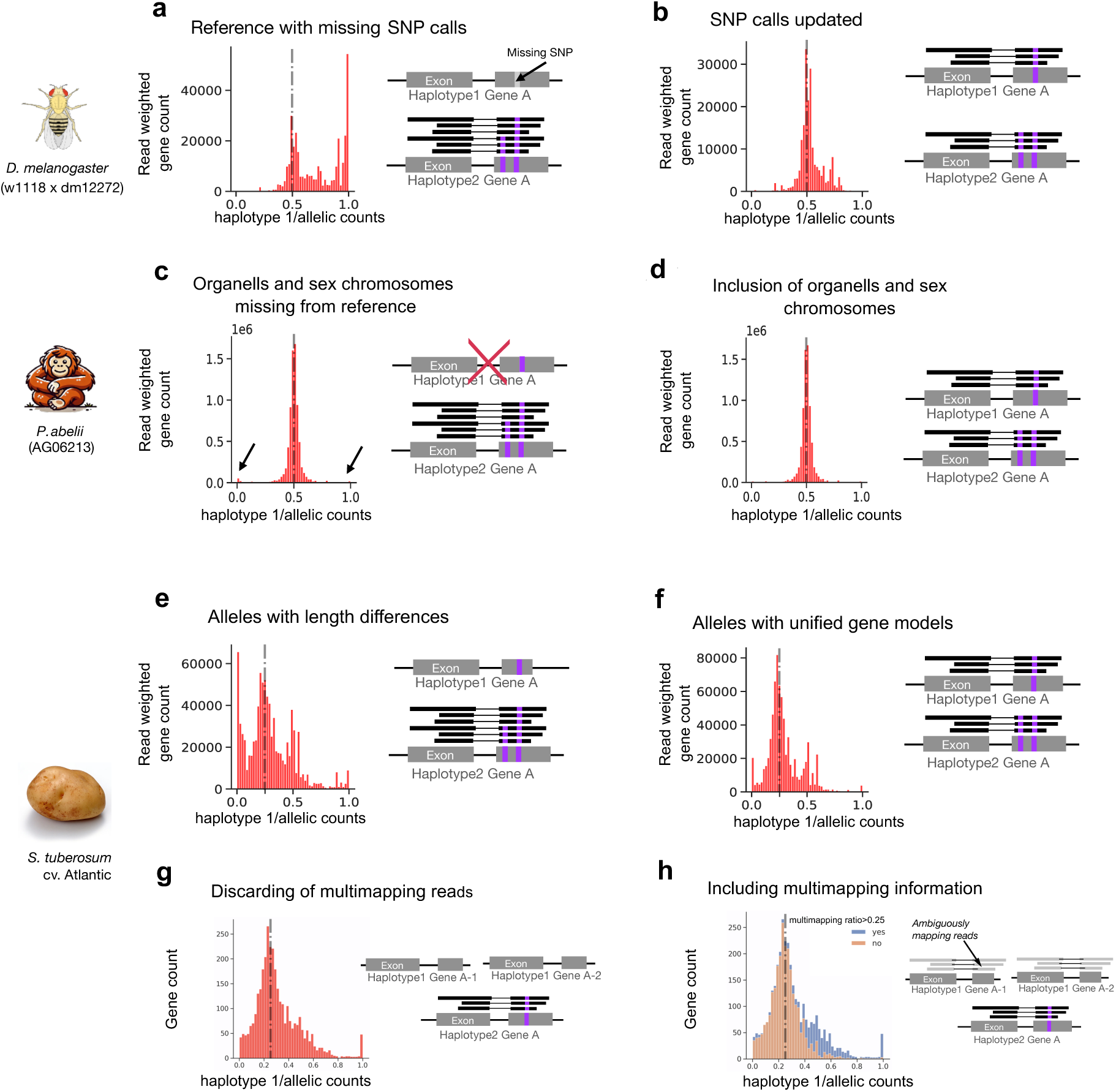
Examples of annotation and assembly errors causing bias that can be identified and corrected with long-read RNA-seq. *(a)* SNP-updated references can introduce bias when there are quality differences in SNP calls between haplotypes. *(b)* Long-read sequencing can address this by correcting for missing SNPs. *(c)* Excluding regions from the reference genome can cause misalignment of reads (arrows), while *(d)* using a complete reference genome resolves such issues. *(e)* Differences in gene lengths between haplotypes can result in reads preferentially aligning to the longer allele, whereas *(f)* unifying annotations across haplotypes using tools like Liftoff reduces this bias. *(g)* Discarding multimapping reads can lead to inaccurate conclusions about allele imbalance, while *(h)* retaining and analyzing multimapping reads provides a straightforward strategy to identify genes affected by ambiguity, such as those with copy number variation. Dashed lines indicate ratios of allelic counts if all alleles have equal expression.

Despite the demonstrated advantage to using personalized references, studies of allele imbalance often fail to incorporate them, perhaps because of the perceived additional bioinformatic effort and the lack of demonstration of the impact of the personalized genome on bias reduction with third generation, long-read, sequencing technology. Long-read RNA-seq studies of allele imbalance (see Supplemental Table S1 for mapping strategies used with long-read RNA-seq) have used two, apparently divergent, mapping strategies. Mapping parameters are sensitive to the complexity of the reference, suggesting that default parameters may need to be adjusted if the reference size and complexity changes, as is expected when transitioning from a haploid to a diploid or polyploid reference or moving from a genome to a transcriptome. While tools and pipelines for allele-specific expression analysis with long-read RNA-seq have been developed, such as LORALS (Glinos et al., 2022), IsoPhase (Wang et al., 2020), IDP-ASE (Deonovic et al., 2017), or IsoLaser (Quinones-Valdez et al., 2024), their use is limited to well annotated and diploid organisms.

We demonstrate using a case study approach of *D. melanogaster*, *Homo sapiens*, *Pongo abelii* and *Solanum tuberosum* how long read RNA-seq can be used improve estimates of allele counts and reduce bias. In addition, we provide a schematic overview of the recommended workflow, summarizing the three main steps and highlighting potential sources of bias and their diagnostic interpretation (Supplemental Fig. S1)

### Recommendation 1: Use a personalized reference

In our first case study, we evaluate allele counts and gene counts mapping to an updated Hg38 compared to mapping to a haplotype specific reference for HG002 using public data. We used ONT cDNA data from three Genome in a Bottle (GIAB) HG002 samples and performed a competitive alignment to the T2T-HG002 assembly and the updated GRCh38 genome (Dwarshuis et al., 2024). We used Liftoff to map the GENCODE v48 annotations onto each haplotype of the HG002 diploid assembly (Shumate & Salzberg, 2021). We classified genes with only a single annotated copy for both haplotypes of HG002 and HGCH38 as singletons and separated those with copy number variation. We quantified, for each gene, the number of reads mapping to both haplotypes in the same gene (Gene-specific GS), reads that mapped to a specific haplotype/allele in one gene (Allele-specific hap1/hap2) using the mapping score *ms* to assign the best alignment(s) for a read. While for most genes the differences were small, there were many genes for which there was a dramatic increase in the ability to assign reads to haplotypes (Supplementary Fig. S2). On average 6.2% more of reads could be assigned allele-specifically in HG002 compared to HGCh38 (Table 1).

**Table 1:** Read assignments based on mapping to the updated GRCH38 or the HG002 assembly. The proportion of mapped reads that are haplotype specific increases for all three samples.

| Sample | Read assignment | GRCh38 (%) | HG002 (%) |
| --- | --- | --- | --- |
| NA27730 | Gene-specific | 77.29 | 71.45 |
| NA26105 |  | 79.27 | 73.31 |
| NA24385 |  | 83.73 | 76.86 |
| NA27730 | Allele-specific (hap1) | 11.16 | 14.78 |
| NA26105 |  | 10.19 | 13.86 |
| NA24385 |  | 8.14 | 12.67 |
| NA27730 | Allele-specific (hap2) | 11.55 | 13.77 |
| NA26105 |  | 10.54 | 12.84 |
| NA24385 |  | 8.13 | 10.47 |

### Recommendation 2: Track multi-mapping reads, do not filter MapQ=0

Although it is often implicitly assumed by individual scientists that the genome size and complexity do not influence mapping, as evidenced by the reflexive use of default parameters, complexity of the reference was recognized to be important by mapping algorithm developers (Kim et al., 2019a; Shumate & Salzberg, 2021). It has been reported that aligning to haplotype-resolved genotype references competitively results in less accurate quantification than aligning in parallel (Hu et al., 2021; T. C. Y. Kuo et al., 2020). One possible explanation for the observation is that default filtering of multi-mapping reads is expected to affect these alignment strategies differently (Fig. 1 c, d).

A transcriptome reference further adds complexity as there will be redundant sequence between isoforms, even in relatively simple transcriptomes such as *D. melanogaster,* and filtering multimapping reads is not common practice when aligning to transcriptomes. The use of equivalency classes, rather than filtering, explicitly acknowledges the uncertainty that arises from the use of the more complex reference. Seesaw (Wu et al., 2023) and Ornaments (Adduri & Kim, 2024) are frameworks that have been designed to quantify short-read RNA-seq data for analysis of allele imbalance by mapping reads with salmon (Patro et al., 2017) or kallisto (Bray et al., 2016) to a diploid-resolved transcriptome and probabilistically assign reads to transcripts and alleles. EMASE uses an expectation maximization algorithm to assign multimappers hierarchically, distinguishing between multi-allele, multi-transcript, and multi-gene reads (Raghupathy et al., 2018). MMSEQ (Turro et al., 2011) is another framework using multimapping reads that was proposed for assigning short-read RNA-seq allele specifically to isoforms using Gibbs sampling.

While probabilistic assignment is an elegant solution, it does not resolve the uncertainty and there is potential for bias in probabilistic assignments (Deschamps-Francoeur et al., 2020; Robert & Watson, 2015; Szabelska-Beresewicz et al., 2023; Zytnicki, 2017). A possible approach to solve this issue is to make use of uncertainty in the quantification output. Alternatively, some researchers have suggested clustering regions with high sequence similarity during gene expression analysis (Robert & Watson, 2015; Sarkar et al., 2020; Zytnicki, 2017) when using transcripts as a reference for mapping (Davidson & Oshlack, 2014; Turro et al., 2014). Simulations can provide insights into the impact of the sequence similarity on the uncertainty and bias (Hafezqorani et al., 2020; L. G. León-Novelo et al., 2014; Lin et al., 2024) and the identification of equivalency classes facilitates the evaluation of the reference and annotation for error.

When mapping short-reads, the high proportion of multimapping in repetitive regions makes tracking multi-mapping reads cumbersome (Fig. 1d). Advances in long-read RNA-seq with the potential to sequence the full transcript, increase the likelihood of capturing single nucleotide and structural variation between haplotypes as the chances of a single read spanning multiple variants are higher (Fig. 1d). We hypothesized that parallel mapping to haplotypes with subsequent read assignment based on the best mapping location and competitive mapping to genotypes should result in comparable estimates of allele specific and gene specific expression assuming that: the accuracy of alignment tools remains unaffected by the reference genome’s size and similarity; that reads multimapping to alleles or loci are identifiable in both approaches; and that the score function is due to the match quality between the read and the reference.

We use the following designations for each read: i) unique mapping to a haplotype/allele, ii) mapping to a gene; or iii) multimapping to multiple genes. We used Minimap2, one of the most popular alignment tools for long read mapping (Li, 2018, 2021). We use this tool here, but the results are general and should not be unduly influenced by this particular choice, although some of the details, as we describe, are particular to the mapping algorithm and we acknowledge that the best alignment may differ based on exactly what function is used. In Minimap2, assignment of allele specific reads can be determined by alignment score (AS), number of mismatches (NM), or primary alignment score (*ms*). A read that maps equally well to more than one location has equivalent alignment scores (AS and *ms* score), however one mapping location will be chosen as primary alignment and the other one(s) as secondary even if the two alignments are identical (in case--secondary=yes and N (number of secondary alignments reported) is > 0). As we wish to compare the sequential and competitive alignment strategies, we opt to use the *ms* score (not AS score) to identify the best match as this setting avoids favoring unspliced mapping to pseudogenes (Li, 2018) (see Supplementary Table S9 for explanation of alignment scores).

### Parallel Mapping to Individual Haplotypes (Fig. 1 c)

This approach aligns reads to the individual haplotypes separately. Allele specific reads are defined by the haplotype with the highest mapping score for each read. Those reads with identical mapping scores to multiple reference haplotypes are combined with the allele specific reads to estimate gene expression (Akama et al., 2014; Glinos et al., 2022; Graze et al., 2012; T. Kuo et al., 2018; L. G. León-Novelo et al., 2014; Munger et al., 2014; Page et al., 2013; Rozowsky et al., 2011). This parallel mapping approach requires an additional script or tool that decides for each read whether the alignment is better to one haplotype or equally good.

### Competitively Mapping to Genotypes (Fig. 1 d)

This approach uses a genotype assembly that consists of a compilation of multiple phased haplotypes (Jallet et al., 2023). The parallel haplotype mapping strategy requires an explicit choice of the criteria to use to consider a read allele specific, with options for alignment scores (As or *ms*), as well as number of mismatches. In contrast, in the competitive genotype mapping strategy, the mapping parameters will be what determines if a read has a primary, allele specific, alignment. Reads that map to multiple haplotypes for a single gene (gene specific reads in the haplotype alignment strategy) will be multi-mapping to the gene in this strategy. Counting the gene specific reads is then by necessity a separate step (Bray et al., 2016; Pandey et al., 2013; Raghupathy et al., 2018; Turro et al., 2011).

We mapped long-read RNA-seq data (PacBio Iso-Seq and ONT) to gene regions for personalized haplotype and genotype references in four species: *D. melanogaster* (fruit fly, Fig. 1e, SNP-updated), *P. abelii* (male Sumatran orangutan fibroblast cell line, Fig. 1f, phased assembly with PacBio Hifi and ONT ultra-long-reads), and *S. tuberosum* cv. Atlantic (potato, Fig. 1g, phased assembly with PacBio HiFi, ONT DNA, Hi-C). Details on these genome assemblies and long-read RNAseq samples are available in Supplemental Table S2. For the four species, we mapped the long-read RNA-seq to the haplotypes in parallel and to the genotypes competitively (see Supplemental Methods for detailed mapping parameters). A read was considered allele specific if the mapping location with the highest *ms* score reported by Minimap2 was unique to one allele. When a read had multiple maximum *ms* scores, it was labeled as “multi-mapping gene specific” if all maximum *ms* scores were observed for the same locus or “multi-mapping multi-gene” when the maximum *ms* score occurred in multiple loci. Tools developed for long-read RNA-seq quantification can be utilized for allele-specific read assignment. For instance oarfish (Zare Jousheghani et al., 2025) assigns reads to transcripts based on *ms* scores and outputs read assignment probabilities and raw unique and ambiguous counts, which informs about the uniqueness and certainty of the alignment.

### Competitive and Parallel mapping strategies are ‘identical’

Mapping to haplotypes and mapping to genotypes was identical for more than 96% of all loci in all the four examples. In the 1.9% - 3.9% of genes with different read counts (Supplemental Table S4), most differ by a single read. Only 0.52% - 0.75% genes have more than 1 read difference between the two mapping strategies. Overall, only 0.021% - 0.033% of all reads have different mapping locations in these two strategies. We hypothesized that these small differences were due to properties of the mapping algorithm and the indexing of the references. Minimap2 is not guaranteed to find the best alignment, the indexing will affect the map location. The impact of the index on the map location could be problematic for large genomes where there are potential areas of sequence similarities, such as in phased haplotypes (Li, 2018). In support of this hypothesis, we observe that for *D. melanogaster* (cumulative size of gene regions on haplotypes: 0.2GB) a smaller percentage of reads had a different alignment position between the sequential haplotype mapping and the competitive genotype mapping compared to the larger gene regions of *P. abelii* (cumulative size of gene regions on haplotypes: 3.0GB; similar to human) and *S. tuberosum* (cumulative size of gene regions on haplotypes: 1.1GB) (Supplemental Table S4).

### Reference complexity affects mapping

Minimap2, like other tools, uses a minimizer index, and parameters for indexing and mapping can influence the primary mapping position found for a read (Kim et al., 2019b; Li, 2018). Genotype references contain more sequence ambiguity, by definition, since multiple alleles are included for each locus. Increasing the threshold for retaining repetitive k-mers can increase the mapping accuracy at cost of longer runtime (see Supplemental Table S4 for the impact of changing default -f 0.0002 to -f 0.000002 for *S. tuberosum*). Using the -a (or -c for .paf output) option in Minimap2 will perform a base-level alignment, which is more accurate than the default reported alignment and we therefore choose to use the base level alignment it in our case study (Li, 2021). Minimap2 with parameter -P (avoids setting of primary chains, see Supplemental methods) resulted in overall higher concordance of the mapping strategies (Supplemental Table S4). To test our hypothesis that mapping parameters affect disagreement between the competitive and parallel mapping strategies, we repeated the case study with the parameter -N 200 (without -P) recommended for transcript quantification (Ji & Pertea, 2024; Zare Jousheghani et al., 2025). As predicted, the discordance between the mapping strategies was slightly higher (e.g loci with different read counts more than 1 read increased from 0.7% to 2.3% *for P. abelii*) (Supplemental Table S4 and Supplemental Fig. S4a-c). In addition, we also observe an increase in concordance of mapping strategies between minimap2 v 2.24 and v 2.28 (Supplemental Table S4).

### Mapping to the transcriptome increases the complexity of the reference

We mapped the reads from one of the *D. melanogaster* genotypes (dm12272) to the haplotype resolved personalized transcriptome using a competitive mapping strategy. We compared mapping to the transcriptome to mapping to the gene regions (described above). As expected, the proportion of multi-mapping reads increased by 10%, reflecting the increase in the equivalency classes in the transcriptome compared to gene regions (Supplementary Figure S5). We examined the impact of mapping to the transcriptome compared to gene-regions on whether the read mapping was allele specific, haplotype specific and gene specific for the *D. melanogaster* data. We found that while there was a large overlap, 77% of the reads were allele specific in both gene region and transcriptome mapping, there were reads that mapped allele specifically to gene regions and not the transcriptome and *vice versa*. The number of reads mapping allele specifically was similar in these two strategies with 1% more reads mapping allele specifically to the transcriptome (Supplementary Table S6). SQANTI3 categories were computed for each read based on genome alignments (Keil et al., 2024; Pardo-Palacios et al., 2024). Reads classified as intergenic, novel-in-catalog, novel-not-in-catalog, antisense and fusion were slightly more likely to map to the gene regions allele specifically than to the transcriptome; while those reads that were classified as full-splice-match, incomplete-splice-match and genic were slightly more likely to map allele specifically to the transcriptome (Supplementary Table S7). This suggests that the discrepancies are again due to the difference in the complexity of the reference and underscores the importance of examining the assumption about the reference complexity in choosing parameters for mapping. Likely this discrepancy could be reduced by further optimization of the mapping parameters.

### Longer reads increase specificity

The average read length for allele specific reads was higher than gene specific and multi-gene multimapping reads. The average read lengths of multimapping reads were 462 bp in *D. melanogaster*, 2031 bp in *P. abelii*, and 748 bp in *S. tuberosum* (Supplemental Table S3). In contrast, the corresponding average read lengths for allele specific reads were 749 bp, 2379 bp, and 838 bp, respectively (Supplemental Table S3). In addition, we observed that for *P. abelii*, only 30% of the reads multimap to the same gene and 68% of the reads mapped allele specifically, whereas in *D. melanogaster* 75% of the reads multimap to the same gene or multiple genes (Supplemental Fig. S3 a,b and Supplemental Table S2). The result is that longer reads reduce the number of equivalency classes.

### Multimapping reads identify genes that share conserved sequences

In *P. abelii* the HOXC4, HOXC6, HOXC8 form a highly conserved gene family. There were 799 reads that multi-mapped to these loci with an average read length of 1786bp. Forty-two reads mapped allele specifically to HOXC4 and none to HOXC6 and HOXC8. In *D. melanogaster* there were 7581 multimapping reads between loci CG31775 and CG42586 and 16 allele specific reads. These two genes are annotated paralogs (Öztürk-Çolak et al., 2024). In both examples, the relatively low number of allele specific reads compared to the high number of multimapping reads, indicates inferences at these genes need to be treated with caution.

### Differences in multi-mapping between haplotypes can signal copy number variation

Differences in annotated structural variation between haplotypes can introduce biases (Van De Geijn et al., 2015). For example, if a gene (or part of a gene) is annotated as duplicated on one haplotype but not the others, multi-mapping reads may appear to reduce the allele specific counts on the haplotype with the duplication. In the *S. tuberosum* the allele on haplotype 4 with a single copy for the gene “translationally controlled tumor protein” (Soltu.Atl_v3.01_4G026870.2, see genomic locus representation in Supplemental Fig. S7a) shows 701 allele specific counts, while other haplotypes’ alleles (Soltu.Atl_v3.01_1G026620.2, Soltu.Atl_v3.01_2G031140.2, Soltu.Atl_v3.01_3G024160.4) show no allele specific counts. However, there are reads multimapping to these loci and their annotated paralogs on haplotypes 1, 2, and 3 (Supplemental Fig. S7a). If multi-locus multimapping reads had been discarded (Fig. 2g), the confounding among these genes would not be clear and an incorrect inference of allele imbalance for “translationally controlled tumor protein” would be reported (Supplemental Fig. S7b). Reporting ratios of multimapping reads vs uniquely mapping reads helps to identify genes with high ambiguity (Fig. 2h).

### Recommendation 3: Use observed extreme bias to identify errors in assembly and annotation

In our case studies of *D. melanogaster*, *P. abelii*, and *S. tuberosum* we demonstrate how to identify and correct errors. In normal tissue, we expect most of the genes to be expressed equally from all alleles. Genes that have a high proportion (e.g. more than 90%) of reads aligning exclusively to one allele, may indicate imprinting or other epigenetic effects, but may also indicate that there are gene annotation or assembly errors in the reference. If the allele imbalance indicates a dominant haplotype this is likely to be due to differences in assembly or annotation quality between the haplotypes rather than reflecting a genuine biological phenomenon (Boatwright et al., 2018; Crowley et al., 2015; Graze et al., 2012; Liu & Wang, 2023). Long-reads make identifying and correcting errors more straightforward (Zhu et al., 2024).

### Missing SNPs

For the *D. melanogaster* experiment the F1 genotypes were a cross where SNPs for the two parental lines relative to the haploid reference genome were called independently for the w1118 line (Campo et al., 2013) and dm12272 line (King et al., 2012). We updated the reference for each of the parents based on the published SNPs and observed an overall bias toward the dm12272 haplotype (Fig. 2a), perhaps not surprising as there were more SNPs in the dm12272 compared to the w1118. Positions predicted to be heterozygous with a dm12272 SNP relative to the reference and w1118 were evaluated. If there was no evidence for the reference allele, we assumed that the SNP was missing from the w1118 calls and included this SNP in the w1118 variant calls. When we remapped using the corrected w1118 variants, haplotype bias was resolved (Fig. 2b).

### Missing genes

Mapping algorithms are “greedy,” meaning they will map reads even with high levels of mismatches. Missing genes in the reference assembly can lead to reads mapping erroneously to other alleles (Fig. 2c). If a gene is absent, reads originating from that gene are likely to map to similar genes elsewhere. This may manifest as allele imbalance if the alleles at the missing locus map preferentially to one of the haplotypes. In the *P. abelii*, the gene *PDHA2* had 1,040 allele specific reads aligned to haplotype 1, none to haplotype 2 and 5 non-allele specifically. This high bias towards haplotype 1 suggested an assembly or annotation error. When the allele specific reads were examined, there was evidence for more than two haplotypes in the alignments suggesting a potential missing gene with similar sequence. The paralog of *PDHA2* known as *PDHA1* is located on the X chromosome. The initial mapping was done for autosomes only and these results suggested that the allele specific reads for *PDHA2* might originate from the *PDHA1* locus. By expanding the reference to include the X and Y chromosomes, as well as mitochondrial DNA, the bias towards *PDHA2* haplotype 1 was resolved. All 1,040 reads previously aligned to the *PDHA2* haplotype 1 allele now aligned to *PDHA*1 on the X chromosome. Indeed, even the 5 multimapping gene specific reads for *PDHA2* aligned to *PDHA1.* In total 1507 genes were located on the X, Y and mitochondrial DNA. When we updated the reference to include these chromosomes, 636,996 reads mapped to these genes and of these 71% aligned in the initial mapping, with 57% of these aligning alleles specifically (145238 to Haplotype 1 and 120488 to haplotype 2) (Supplemental Table S5). Including the missing chromosomes decreased the number of loci with apparent extreme bias (Fig. 2d).

### Length variation between haplotypes

Incorrect exon–intron structures are common artifacts of genome annotations (Y. Chen et al., 2023; Mathé & Dunand, 2021; Meyer et al., 2020; Salzberg, 2019). In the *S. tuberosum* assembly the gene annotation was performed per haplotype, and genes were not explicitly matched between haplotypes during annotation (Hoopes et al., 2022). Alleles with more than a 10% difference between the maximum and minimum lengths of annotated genes showed higher proportions of allele specific reads mapping to the longest allele (Fig. 2e). For 5,099 of the 9,130 genes with all four alleles present in the assembly, there was more than a 20% difference in the length of the spliced representative transcript (Supplemental Fig. S6a). We unified the annotation between haplotypes using Liftoff (Shumate & Salzberg, 2021), which led to more similar transcript lengths between haplotypes (Supplemental Fig. S6b) and more balanced allelic ratios (Fig. 2f). For example, the number of genes with allelic ratio > 0.9 towards haplotype 1 decreased from 136 to 61. A reduction in extreme bias was observed for all haplotypes (Supplemental Fig. S8). Long-read RNA-seq can also be used to improve existing annotations by identifying unannotated transcripts and genes with tools like BAMBU (Y. Chen et al., 2023). But to avoid bias due to read misassignment it needs to be ensured that the same transcript models are identified on all haplotypes.

### Conclusion

Haplotype-resolved telomere-to-telomere genomes are available for diploid individuals (Han et al., 2023; Yang et al., 2023) to highly polyploid plant cultivars (Healey et al., 2024; Zheng et al., 2022). The next step is the construction of pangenomes. Pangenomes incorporate population level genetic variation (N. Chen et al., 2021; Coombes et al., 2024; Sibbesen et al., 2023; Yuan et al., 2015) and tools for pangenome mapping (Chandra et al., 2024; Sibbesen et al., 2023) and allele identification and expression estimates for human pangenomes are being developed.

Allelic variation, copy number variation, ploidy and gene families all contribute to sequence similarity and lead to uncertainty. The degree of uncertainty will vary across the genome and how best to identify and incorporate uncertainty into estimates of allelic and gene expression will take time and effort. Meanwhile, the practice of filtering multi-mapping reads should be discontinued, even when aligning in parallel to haplotypes. Although read lengths increase specificity of mapping, not all reads will be allele, or gene specific. In our case studies, 30 – 75% of reads were gene specific, but not allele specific, and 2-9% of reads multimapped between loci. Tracking multimapping reads for reads that mapped to multiple loci identified uncertainty and enabled correcting mis-assemblies and mis-annotations and the identification of hidden paralogs. Tracking reads that differ in location, and number of alignments between haplotypes, may also pinpoint hidden polymorphisms in copy number variation (CNV) between the haplotypes. CNVs are difficult to disentangle and especially challenging when they are recent expansions.

These considerations are particularly relevant in the context of cancer, where somatic structural variation, aneuploidy, and copy number changes make allele-specific analyses especially challenging (Baker et al., 2024; Cortés-Ciriano et al., 2020; Drews et al., 2022; Garribba et al., 2023; Klockner & Campbell, 2024; The Cancer Genome Atlas Network, 2015). Improved assemblies and haplotype-resolved references for cancer genomes would substantially enhance our ability to distinguish true regulatory differences from artifacts introduced by structural complexity. This would have direct implications both for clinical diagnostics, where accurate detection of allelic imbalance may inform patient stratification, and for research projects aimed at understanding tumor evolution and regulatory mechanisms in cancer.

The success of long reads in resolving complex structural polymorphisms in DNA (Chang et al., 2022; Courret & Larracuente, 2023; Logsdon et al., 2024; Porubsky & Eichler, 2024) presents an opportunity to re-examine pipelines for allele imbalance with long-read RNA-seq. In our case studies we show how personalized references and tracking multimapping of the long-read RNA-seq data can be used to identify, and correct mis-assembly and mis-annotations using relatively straightforward bioinformatic techniques and existing tools, counter to the perception that using personalized reference, and tracking multi-mapping reads is complicated and time consuming. Although tools for allele-specific long-read RNA analysis exist, there is currently no universally applicable workflow that can be used for different organisms. Our recommendations apply to diploid and polyploid organisms as well as model and non-model organisms.

## Code availability

Nextflow (Di Tommaso et al., 2017) pipeline for allele specific mapping to haplotype and genotype references, mapping comparison and plots is available at https://github.com/nadjano/nf-ASE-mapping-comparison.

Additional scripts for reference preparation are available at https://github.com/nadjano/nf-ASE-mapping-comparison/tree/main/scripts_for_reference_preperation.

The lifoff gene annotations for S. tuberosum cv Atlantic and H. sapiens HG002 have been deposited to <u>Zenodo</u>.

## Declarations

LM has received reimbursement of travel and accommodation expenses to speak at Oxford Nanopore Technologies (ONT) conferences.

All other authors declare no competing interests.

## Acknowledgements

We thank the Eichler Lab for their work on the Pongo abelii phased assembly and sharing the gene Liftoff annotation and Alison Morse for work on the Drosophila data

This work has received funding from the European Union’s Horizon 2020 research and innovation programme under the Marie Skłodowska-Curie Actions Doctoral Network “LongTREC” grant agreement No 101072892; and the Slovenian Research and Innovation Agency grant agreements no. P4–0165 and P4–0431; Associazione Italiana per la Ricerca sul Cancro (AIRC) (IG22851), National Center for Gene Therapy and Drugs based on RNA Technology (CN00000041) supported by European Union – Next Generation EU, Mission 4, Component 2, CUP B93D21010860004; Ministry of Health – bando POS – tr.3 – T3-AN-04 – GENERA; an EMBL ETPOD Fellowship; the NIH funding GM137430 and the department of Molecular Genetics and Microbiology, The University of Florida Genetics Institute, the University of Florida Cancer Center, and the University of Florida Research Computing Center (www.rc.ufl.edu). FN is a member of the Human RNome Consortium.

Thanks to all the members of the LongTREC network for discussion.

## Data Availability Statement

The long-read raw data for our case study analyses are available at NCBI’s Sequence Read Archive under accessions SRR29784420-SRR29784421 (*D. melanogaster*), SRR27438208 (*Pongo abelii*), and SRR14993892-SRR14993894 (*Solanum tuberosum* cv Atlantic). For *H. sapiens* NA24385, NA27730, NA26105 from the Genome in a Bottle HG002 ONT cDNA project were used.

## Supplemental Material

### Supplemental Methods

*Supplemental Tables*

Supplemental Table S1: Examples of studies using long-read RNA-seq for quantification of allele specific expression analysis.

Supplemental Table S2: Details about fasta, gff and long-read RNA-seq for each case study.

Supplemental Table S3: Mapping statistics for long-read RNA-seq from *D. melanogaster*, *Pongo abelii* and *Solanum tuberosum*.

Supplemental Table S4: Mapping statistics for different minimap2 parameters.

Supplemental Table S5: Number of reads that align elsewhere in the genome when Chromosome X, Y and mitochondrial DNA are absent from the reference.

Supplemental Table S6: Comparison of read assignment category of reads mapped to transcriptome vs gene regions

Supplemental Table S7: Comparison of read assignment and sqanti3 category of reads mapped to transcriptome vs gene regions

Supplemental Table S8: Description of alignment score metrics used by minimap2

*Supplemental Figures*

Supplemental Figure S1: Flow chart of recommended steps and tools for allele-specific expression analysis.

Supplemental Figure S2: Agreement plot of allele and gene counts between using GRCh38 and HG002 as a reference genome.

Supplemental Figure S3: Barplot of number of reads per mapping category for reads that have identical mapping position in competitive and separate mapping.

Supplemental Figure S4: Heatmaps of read number per category of reads having different mapping locations in competitive and separate mapping.

Supplemental Figure S5: Barplot of length differences between the alleles of genes of the representative annotated transcripts for *Solanum tuberosum* based on the gff file.

Supplemental Figure S6: Example of Copy number variation in *Solanum tuberosum* haplotypes.

Supplemental Figure S7: Distribution of allelic ratios per genes and haplotype for the different length difference categories before length correction after length correction

