## Supplemental Figures for "The promise of long-read RNA-seq: reducing bias in analyses of allele imbalance"

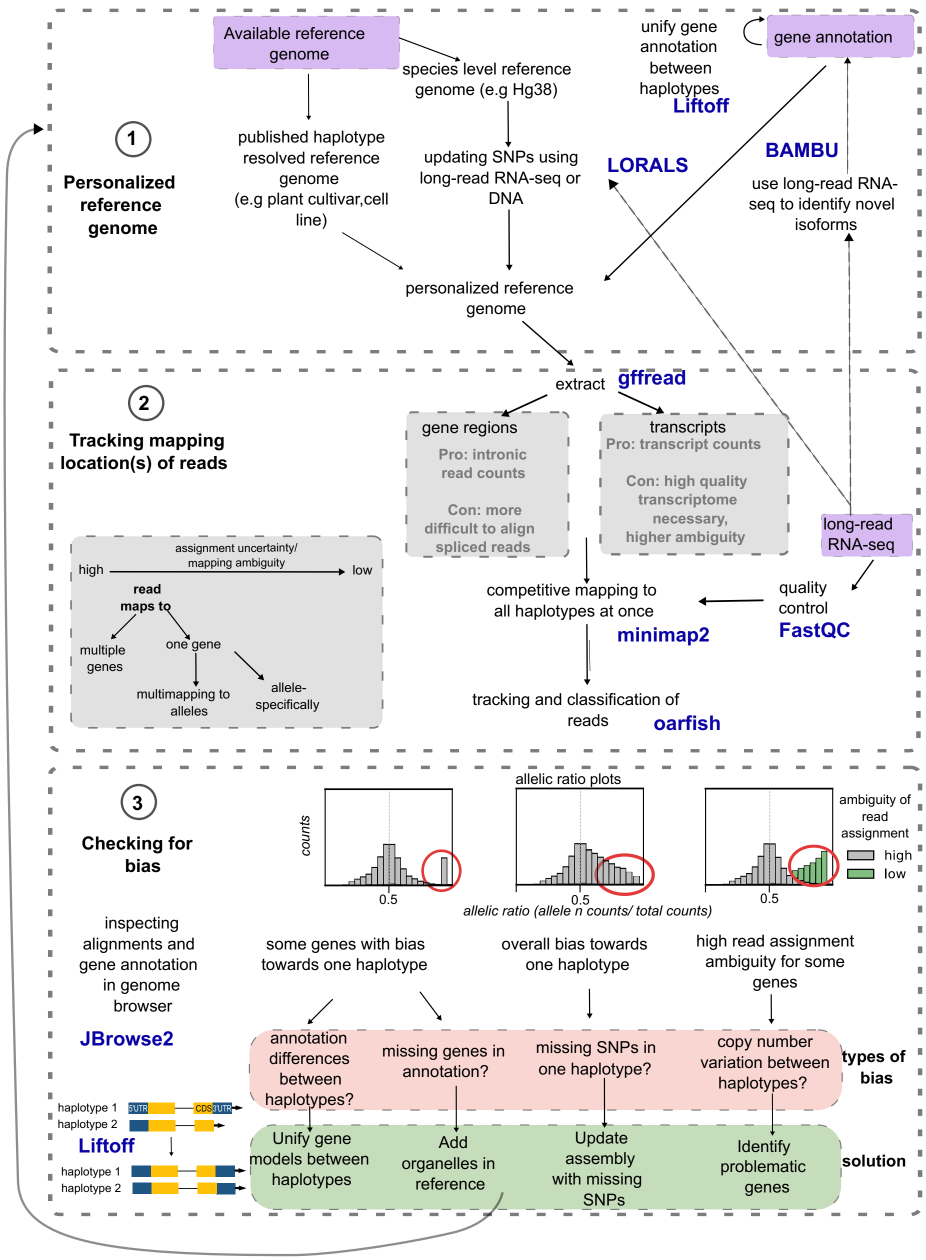


**Figure S1: Schematic overview of the recommended workflow for allele-specific expression analysis using long-read RNA-seq.** The flowchart is organized in three main steps: (1) construction of a personalized reference genome from available assemblies or haplotype-resolved sequences; (2) tracking the mapping locations of reads, including explicit classification of multi-mapping events; and (3) evaluating apparent bias as a diagnostic tool to identify assembly or annotation errors. This figure is intended as a practical guide: each panel corresponds to a decision point in the analysis, and references to the main text indicate where examples and case studies are described. Recommended tools for each step are highlighted in blue.

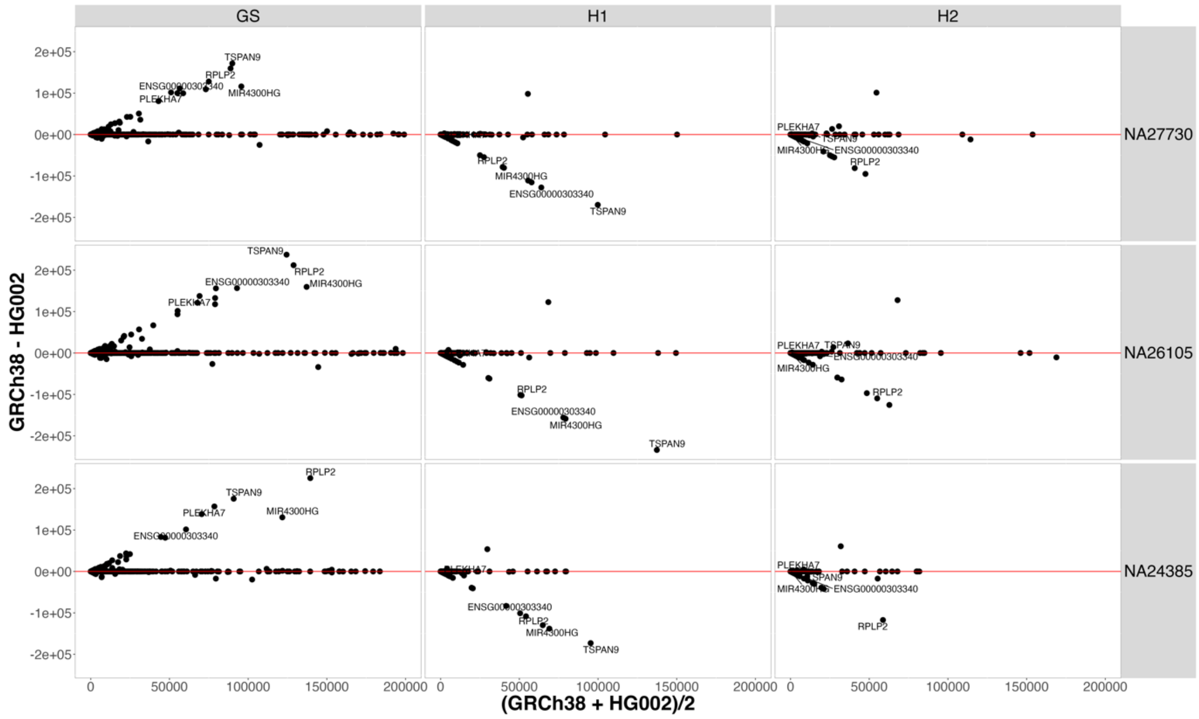


**Figure S2:** Agreement plot showing, for each sample, the difference in counts between the two assemblies (GRCh38 and HG002) (y-axis) versus the mean (x-axis) for each gene. The red line is at y = 0, indicating no difference in quantification between assemblies. The gene names of the top five genes with the largest differences are shown.

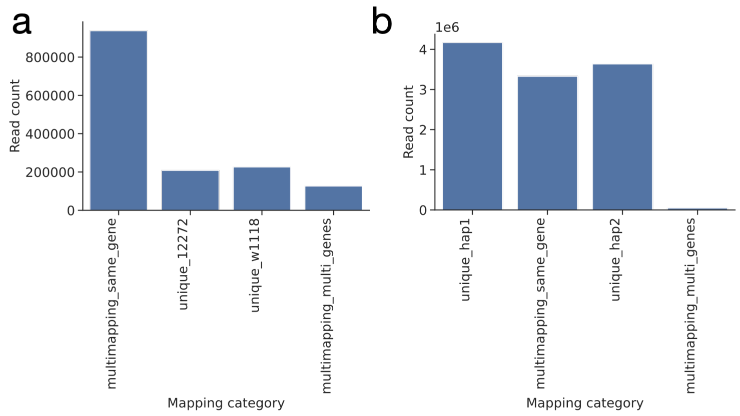


**Figure S3:** Barplot of number of reads per mapping category for reads that have identical mapping position in competitive and separate mapping for a) *D. melanogaster* and b) *P. abelii*. Note: For *P. abelli* Chromosome X, Y and mitochondrial DNA are included in haplotype 1.

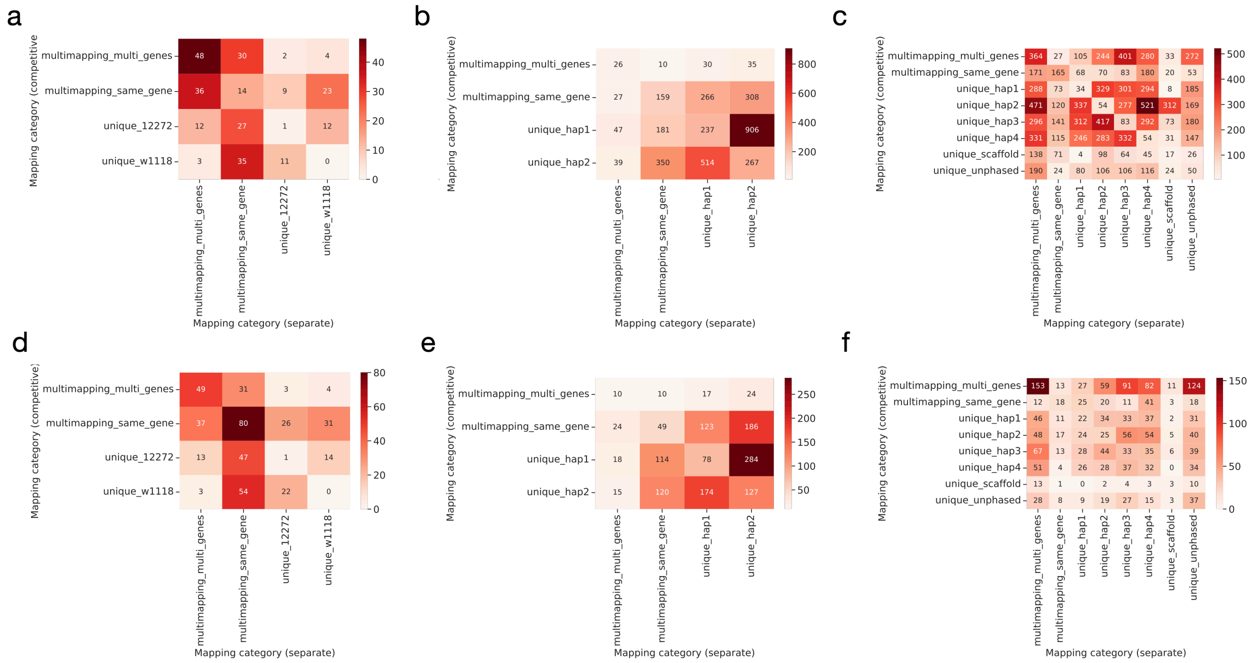


**Figure S4:** Heatmaps of read number per category of reads having different mapping locations in competitive and separate mapping for running minimap2 v2.28 with parameter **-N 200** a) *D. melanogaster* and b) *P. abelii* and *c) S. tuberosum*, and parameter **-P** d) *D. melanogaster* and e) *P. abelii* and *f) S. tuberosum.*

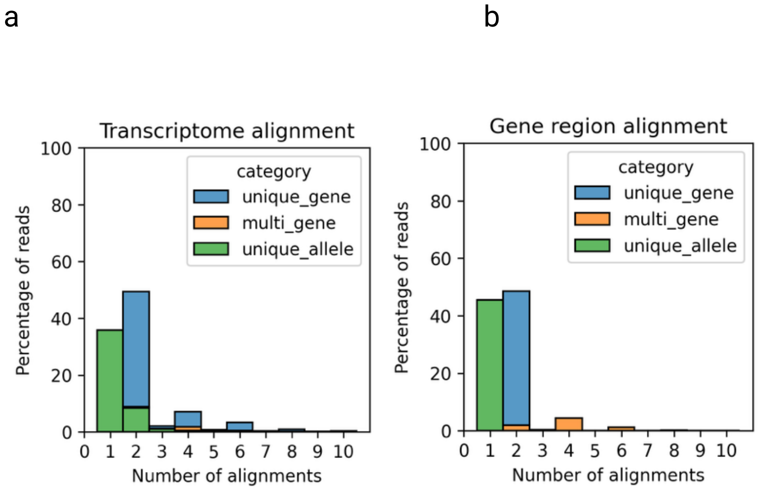


**Figure S5:** Barplots of number of alignments per read (x-axis) in percent (y-axis) and their category of read assignment for a) transcriptome and b) gene region alignments.

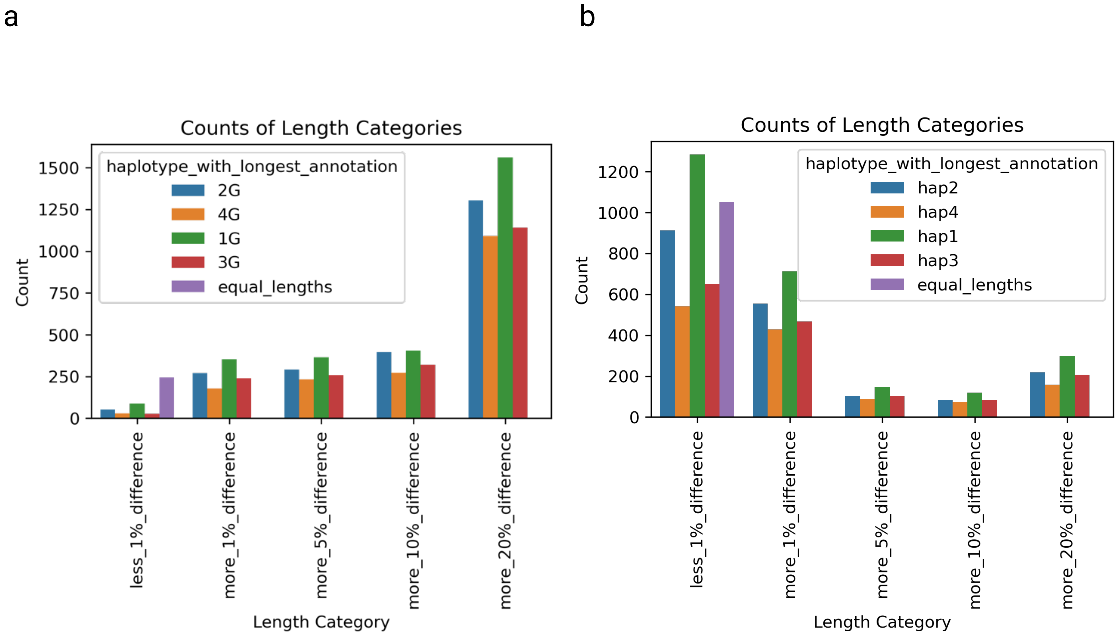


**Figure S6:** Barplot of length differences between the alleles of genes of the representative annotated transcripts for *S. tuberosum* cv Atlantic based on a) the published gff file and b) after correction of gene models with liftoff.

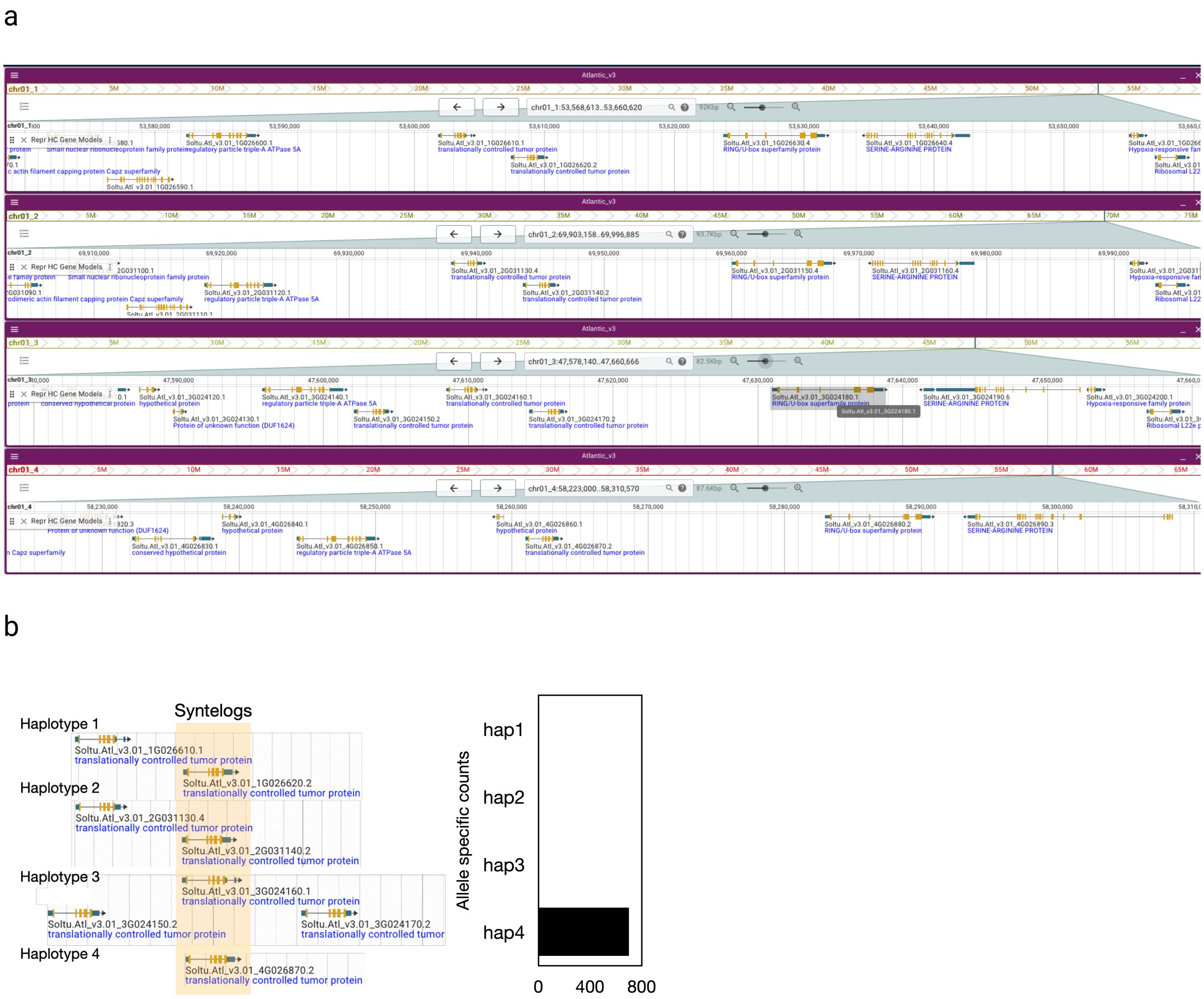


**Figure S7:** Copy number variation in *S. tuberosum* cv Atlantic. a) Screenshot from jbrowse spuddb http://spuddb.uga.edu/atl_v3_jb2/?session=share-NBYY55MwNL&password=700KB. b) Visualization of copy number of "translationally controlled tumor protein" per haplotype and allele-specific counts for the "translationally controlled tumor protein" per haplotype.

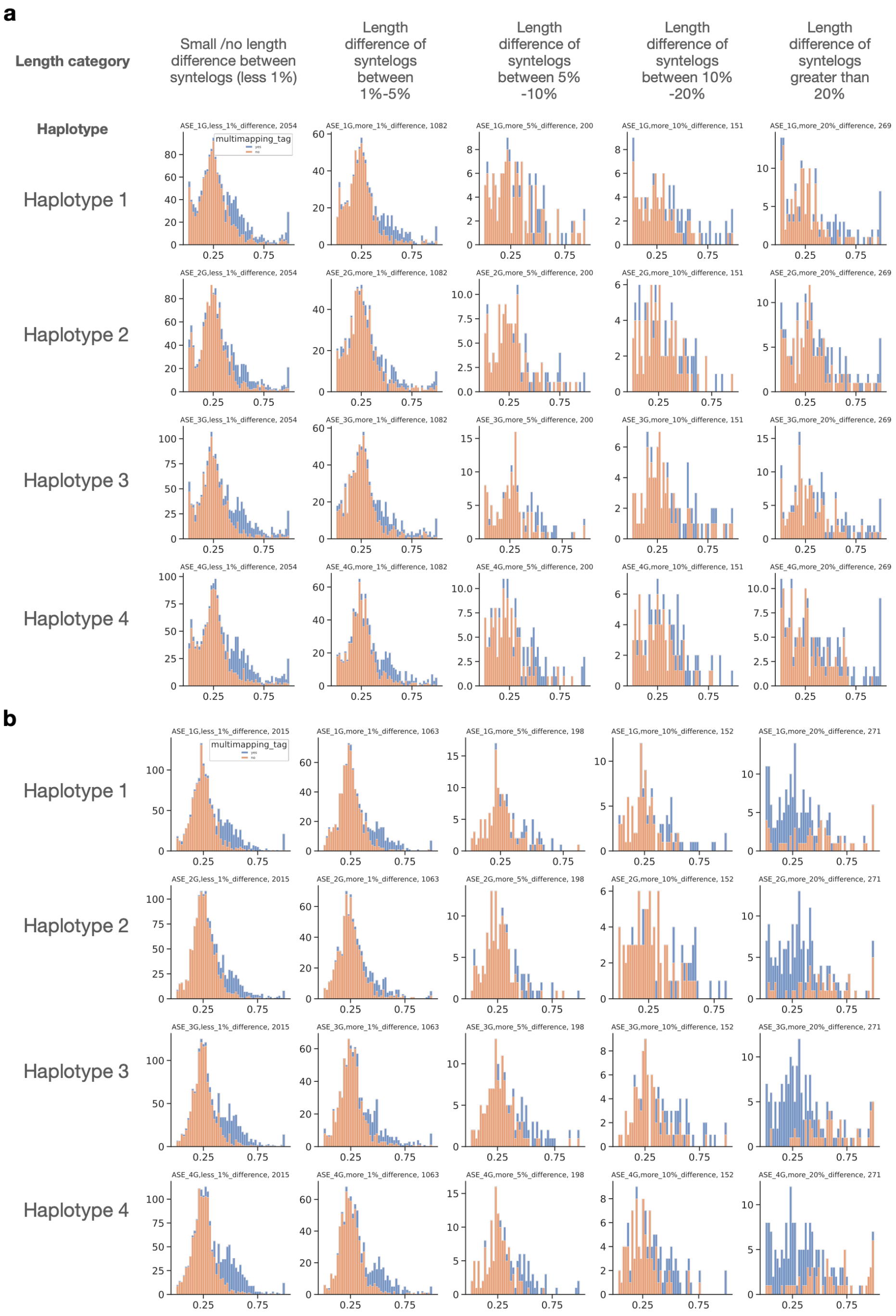


**Figure S8**: Histograms showing distribution of allelic ratios per genes and all four haplotypes for the different length difference categories a) before length correction b) after length correction. X-axis show ratio of reads per gene that align allele-specifically. Y-axis show number of genes. Plot title indicates haplotype and length differences between syntelogous genes. Colors represent fraction of multimapping reads per gene. Orange: less than 25% of total reads are multimapping.

Blue: more or equal 25% of total reads are multimapping.
