## Supplemental Methods for "The promise of long-read RNA-seq: reducing bias in analyses of allele imbalance"

**Supplementary Methods**

**Preparation of references**

The dmel6 reference genome was updated with w1118 and dm12272 specific SNPs respectively. SNP positions where updated, in dmel6 to produce to different genomes.

To correct for observed allelic bias to the dm12272 genotype, long-reads were aligned and positions with SNPs (min reads 5) in dm12272 and no SNPs in w1118 but with no reads only containing a SNP were added to the w1118 VCF file and gene regions were extracted again.

For *H. sapiens*, the T2T genome assembly of HG002 cell line by GIAB and the GRCh38 genome were used. Reference genomes were updated with phased SNPs (GIAB’s High-confidence variant calls set). To generate the GRCh38 genome updated with phased SNPs, the phased SNPs were extracted from the HG002 high-confidence variant calls vcf file (Supplemental Table S2).

bcftools consensus was used to generate a fasta file for GRCh38 and the two haplotypes of HG002, with haplotype 1 being the paternal (containing chrY) and haplotype 2 the maternal (containing chrX and chrM). Additional non-chromosomal contigs were removed. Next, we annotated both haplotypes using Liftoff v1.6.3 (Shumate & Salzberg, 2021) and the GENCODE v48 annotation on the GRCh38 genome, applying the -polish and -sc 0.9 options to search for additional gene copies with at least 90% identity. Genes were annotated as singletons if they had a single copy in all haplotypes and assemblies, the difference in length (counting the sum of exon lengths) did not exceed 25 bp, they were on the same chromosome in all haplotypes and assemblies, and they had the same number of exons. Gene regions were then generated from the annotations using bedtools getfasta.

For *D. melanogaster*, *H. sapiens*, *P. abelii* and S. tuberosum cv Atlantic gene regions (annotated mRNA +- 150) were extract from the reference genome for all the haplotypes with the respective gff files (see supplementary Table S2). For *S. tuberosum* cv Atlantic the unphased haplotype (hap0) and scaffold s and for *P. abelii* the X and Y chromosomes on haplotye1 and mitochondrial DNA were included. Sequences were extracted with GffRead (Pertea & Pertea, 2020) .

**Minimap2 parameters for comparing parallel and competitive alignment:**

Long-read RNA-seq data were aligned using minimap2 (v 2.12-r827) (Li, 2021) to haplotype-specific FASTA files containing gene regions, either individually or concatenated (e.g., D. melanogaster: w1118, 12272, *S. tuberosum* cv Atlantic: hap1, hap2, hap3, hap4, hap0 (unphased); *P. abelii*: hap1, hap2) using parameters -c -x splice --secondary=yes -P. The parameter P was chosen to find all alignments and not set primary alignments, as we observed this led to a higher agreement between the sequential mapping to haplotypes and whole genotype mapping than running with parameter -N 200 (Supplemental Table S4). For quantification, it is recommended to increase the -N parameter (maximum number of secondary alignments output) from the default of 5 to 100 (Jousheghani & Patro, 2024) or 181 (Ji & Pertea, 2024) to capture all secondary alignments.

For *H. sapiens,* competitive mapping was performed with minimap2 v1.28 using the parameters -ax splice --secondary=yes -N 200.

A custom Python script was used to extract, for each read, the alignments with the maximum mapping score (*ms*) and to classify each read as Haplotype1-specific or Haplotye2-specific (aligned to a single gene on a single haplotype), gene-specific (GS; aligning to both haplotypes), or multigene–multimapping. For each gene we quantified the number of haplotype 1, haplotype 2 and gene-specific reads.

**Read counting**

Reads per gene were counted with custom python scripts available in a nextflow pipeline https://github.com/nadjano/nf-ASE-mapping-comparison/tree/main

**Transcriptome alignment and counting**

To compare the difference of using the transcriptome instead of gene regions. Samples from D. melanogaster were aligned to SNP updated gene regions and transcriptome.

Minimap2 parameters for gene regions: -ax splice -t 16 --secondary=yes -N 200

For transcriptome: -a splice -t 16 --secondary=yes -N 200.

Oarfish (0.8.1) (Zare Jousheghani et al., 2025) was utilized for read assignments to transcripts/gene regions with the parameters: --write-assignment-probs --filter-group no-filters -s 1 –verbose. The parameters -s 1 only considered secondary alignments that have equal *ms* score as primary alignments. The resulting read assignment probability files where parsed and reads where assigned to categories based on their best alignment(s). 1) read aligns best to one transcript 2) read aligns best to two transcripts of the same gene and haplotype 3) read aligns to two transcript, same gene and different haplotypes. To get gene level counts, transcript counts per gene were summed.

Ji, H. J., & Pertea, M. (2024). *Enhancing transcriptome expression quantification through accurate assignment of long RNA sequencing reads with TranSigner*. https://doi.org/10.1101/2024.04.13.589356

Jousheghani, Z. Z., & Patro, R. (2024). *Oarfish: Enhanced probabilistic modeling leads to improved accuracy in long read transcriptome quantification*. https://doi.org/10.1101/2024.02.28.582591

Li, H. (2021). New strategies to improve minimap2 alignment accuracy. *Bioinformatics*, *37*(23), 4572–4574. https://doi.org/10.1093/bioinformatics/btab705

Pertea, G., & Pertea, M. (2020). GFF Utilities: GffRead and GffCompare. *F1000Research*, *9*, 304. https://doi.org/10.12688/f1000research.23297.2

Shumate, A., & Salzberg, S. L. (2021). Liftoff: Accurate mapping of gene annotations. *Bioinformatics*, *37*(12), 1639–1643. https://doi.org/10.1093/bioinformatics/btaa1016

Zare Jousheghani, Z., Singh, N. P., & Patro, R. (2025). Oarfish: Enhanced probabilistic modeling leads to improved accuracy in long read transcriptome quantification. *Bioinformatics*, *41*(Supplement_1), i304–i313. https://doi.org/10.1093/bioinformatics/btaf240
