## Supplemental Table S1 for "The promise of long-read RNA-seq: reducing bias in analyses of allele imbalance"

**Supplementary Table 1:** Overview of studies using long-read RNA-seq for quantification of allele-specific expression analysis.

|  | **Study** | **Species** | **Reference input** | **Mapping strategy** | **Assignment of reads to haplotypes** |
| --- | --- | --- | --- | --- | --- |
| **Sequential haplotype mapping** | (Glinos et al., 2022) | human | Diploid genome, long-read RNA-seq was used to correct haplotypes | Mapping to haplotypes **sequentially** | Counting the number of read containing REF or ATL allele |
|  | (Zheng et al., 2022) | Hexaploid bamboo | Reference genomes from parents | Aligning to parental genomes with minimap2 **sequentially** | Assignment of reads based on SNPs |
|  | (Feng et al., 2021) | Hybrid rice | Reference genomes from both parents | Aligning to parental genomes with minimap2 **sequentially** | Assignment of reads based on SNPs |
| **Competitive genotype mapping** | (Hughes et al., 2023) | Human, HLA | Human reference genome, screening HLA transcript database with long-reads and generation of reference with allele-specific HLA types. | Aligning to HLA type personalized reference genome **competitively** with minimap2 | Assignment based on primary minimap2 mapping. |
|  | (Bresnahan et al., 2023) | reciprocal honeybee crosses | combined parental transcriptomes | Alignment with minimap2 **competitively** to transcriptomes | Not clear: potentially assignment based on primary minimap2 mapping. |
|  | (Dominguez-Ortiz et al., 2024) | human ovarian cancer | annotated Gencode. v44 BRCA1 transcripts | alignment with salmon with long-read parameters **competitively** | Not clear: potentially assignment based on primary mapping. |
|  | (Cornaby et al., 2022) | human HLA | Human reference genome, screening HLA transcript database with long-reads and generation of reference with allele-specific HLA types. | Aligning to HLA type personalized reference genome **competitively** with minimap2 | Assignment based on primary minimap2 mapping. |
| **Mapping to a single haplotype** | (Park & Cenik, 2024) | human and mouse | **N-masked** reference genome | alignment with minimap2 to reference transcriptome | Assignment based on known SNPs |
|  | (S.-Y. Chen, 2022) | F1 hybrids of cattle and yak | **N-masked** reference genome | Alignment with minimap2 | Assignment based on distinguishable interspecies SNPs |
|  | (Xu et al., 2024) | Hybrid maize (B73 × Ki11, Ki11 × B73) | maize RefGen_v4 | Aligning to one parent with minimap2 | **Phasing** of long-read RNA-seq reads over SNP positions with IsoPhase (Wang et al., 2020) |
|  | (Low et al., 2020) | Angus and Brahman crossbred cattle | Reference genome from one parent | Aligning to one haplotype with minimap2 | **Phasing** of long-read RNA-seq reads over SNP positions with IsoPhase (Wang et al., 2020) |
|  | (Wang et al., 2020) | Hybrid maize (B73 × Ki11, Ki11 × B73) | Reference genome from one parent | Aligning to one haplotype with minimap2 | **Phasing** of long-read RNA-seq reads over SNP positions with IsoPhase (Wang et al., 2020) |
